# Host lifestyle and parasite interspecific facilitation mediate co-infection in a species-poor host-parasite system

**DOI:** 10.1101/2023.01.03.522369

**Authors:** Nikol Kmentová, Armando J. Cruz-Laufer, Leona J. M. Milec, Tanisha Moons, Senne Heeren, Elze van den Hoorn, Kelly J. M. Thys, Lawrence Makasa, Auguste Chocha Manda, Pascal Masilya Mulungula, Maarten Van Steenberge, Michiel W. P. Jorissen, Maarten P. M. Vanhove

## Abstract

Despite their important ecological role, questions remain on mechanisms structuring parasite assemblages. We present a simple and endemic host-parasite system of clupeid fishes and monogenean parasites (*Kapentagyrus*, Dactylogyridae) with contrasting levels of host-specificity from Lake Tanganyika as a model to study parasite distribution patterns and co-infection dynamics in nature. With two parasites, two host species, and three host-parasite combinations between them, this unique system represents the simplest natural host-parasite model that is not trivial. We modelled spatiotemporal dynamics of host-parasite interaction using infection data along the North-South axis of Lake Tanganyika (660 km) over the course of two seasons and four years (1730 fish, 3710 parasites). We found temporal stability of infection, which contrasts with previously reported seasonally driven fluctuations of fish host abundances. We found a difference in spatial structure between the parasite species, confirming that their distributions are only restricted by their most mobile host species. On the host species that is infected by two parasite species, we discovered a positive correlation with host body size for one parasite species, and a negative correlation for the other species. As we also discovered facilitation of infection, this cannot be due to competition. The differences reported between parasite species infecting the same host species further extrapolate the dependence on changes in lifestyle of the host during its ontogenetic development. In conclusion, we show that in a simple, closed system parasite infection dynamics are dependent on a combination of host mobility, host lifestyle changes over ontogenetic development and interspecific interactions between parasites.

## Introduction

### Questions remain on mechanisms structuring parasite assemblages

Metazoan parasites are a major part of global species diversity (Poulin, 2014; Windsor, 1998). Despite a largely negative perception, parasites have been recognised as ecosystem engineers that form a substantial biomass in aquatic ecosystems (Kuris et al. 2008) and that alter food web topography, including competition and predation, through their indirect effect on host abundance (Hatcher et al. 2012). They contribute to the energy transfer between trophic levels in ecosystems (Lafferty et al. 2006), e.g. by inducing changes in host behaviour that influence predation (Lefèvre et al. 2009), or by acting as mediators in biological invasions (Blackburn et al. 2011). Metazoan parasites of fish have been studied in terms of general principles shaping parasite population dynamics and community structure at both local (Bagge et al. 2004) and global scales (Morand et al. 1999). The discovery of recurring patterns is key towards understanding general principles in ecology. Through repeated empirical studies across various host-parasite systems, general laws underlying the structure of parasite communities and populations are recognised (reviewed in Poulin 2007). These include a positive relationship between parasite species richness and/or intensity of infection and host body size (Morand et al. 1999, Šimková et al. 2000, Henriksen et al. 2023), the aggregation of parasite populations among host individuals (Morand et al. 1999, Poulin 2013), and more recently the observation that gene flow in a parasite species is determined by its most mobile host species (Criscione and Blouin 2004, Blasco-Costa et al. 2012, Kmentová et al. 2021). Several other patterns characterising infection dynamics of parasite populations and communities have been suggested but lack repeatability across time and space (Poulin 2007), for example the link between host species richness and parasite population size (Bagge et al. 2004), and the effect on parasite infection dynamics of host density (Cote and Poulin 1995, Stumbo et al. 2012, Buck and Lutterschmidt 2017), geographic distance (Poulin and Morand 1999, Poulin 2003, Timi et al. 2010) and interspecific interactions among parasite species (El Hafidi et al. 1998, Kennedy 2001, Salgado-Maldonado et al. 2019, 2020). The ecological study of natural host-parasite systems is often hampered by confounding effects and complexity (Poulin 2007). The confounding effects of host evolutionary relationships (Luque et al. 2004, Sasal et al. 2008), seasonal dynamics (Poulin 2020) and parasite life-cycle complexity (Lima et al. 2012, Wood et al. 2014) influence conclusions on mechanisms driving parasite assemblages. Additionally, since parasite diversity is understudied and large-scale monitoring is lacking, studies underlying processes along the entire distribution range of the respective host and parasite species are rare.

### Lake Tanganyika, a natural laboratory

Lake Tanganyika (LT) has been proposed as a natural study system due to its ecological, geographical and evolutionary delimitation and the negligible anthropogenic impact compared to other aquatic ecosystems worldwide (Coulter, 1991a; Cristescu et al., 2010). While the littoral zone is species-rich in terms of both fish hosts and parasites, the pelagic community is generally species-depauperate and, in contrast to the marine open water realm, clearly delineated within the lake’s confines (Paugy and Lévêque 2017). Pelagic fish hosts are often described as parasite species-poor compared to littoral host communities (Marcogliese 2002). Hence, pelagic host-parasite communities offer a simplified study system to address questions on infection dynamics. The pelagic zone of LT is dominated by two clupeid species, *Limnothrissa miodon* (Lake Tanganyika sardine) and *Stolothrissa tanganicae* (Lake Tanganyika sprat) (Mannini et al. 1996). Clupeid fishes in LT are parasitised by two gill-infecting species of *Kapentagyrus* (Kmentová et al., 2018). *Kapentagyrus* is a genus of monogenean flatworms (Monogenea, Dactylogyridae) whose representatives have so far only been reported in African freshwater clupeid hosts (Vanhove et al., 2021). While *K. tanganicanus* parasitises both clupeid host species, *K. limnotrissae* is specific to *L. miodon*. Many small pelagic fishes, including sardines and other clupeids, are known for their schooling behaviour and large distribution ranges linked to their long-distance migrations (Teske et al. 2021). Given their economic importance to fisheries, spatio-temporal dynamics of both clupeid hosts are well studied. Seasonally-dependent algal blooms (May-June in the South of LT, October-November in the North of LT) driven by upwelling of nutrient rich waters (Plisnier et al. 1999) promote geographical migration of these clupeids (Phiri and Shirakihara 1999, van Zwieten et al. 2002). Recent genomic studies confirmed previously suggested restricted migration of *L. miodon* compared to *S. tanganicae* along the entire North-South axis of LT (De Keyzer et al. 2019, Junker et al. 2020). These clupeid fishes and their monogenean parasites provide a simple model to study spatio-temporal parasite infection dynamics given the low species richness, contrasting host-range, absence of other gill metazoan parasites (Kmentová 2019), sister-group relationship of both fish hosts (Wilson et al. 2008, Milec et al. 2022) and parasite species (Kmentová et al. 2022), and geographically restricted natural distribution of both hosts and parasites (endemic to LT). A large evolutionary distance to other fish (Wilson et al. 2008, Milec et al. 2022) and monogenean taxa present in the lake (Kmentová et al. 2022) further restricts additional host-parasite interactions as confounding factors.

In this study, we investigate spatio-temporal distribution and co-infection dynamics of parasites using a simple host-parasite model consisting of two parasite species infecting two species of clupeid hosts in three host-parasite combinations. We survey parasite infection over the entire natural distribution range of all species involved (Fig. 1). We combine results with data on size and morphology of the fish hosts and with previously published information on the biology and the genetic population structure of the monogenean parasites and their clupeid hosts (Fig. 1).

We hypothesise that:

1. given the host-range difference between the two parasite species and the restricted migration of one of the hosts, *L. miodon* (Junker et al. 2020), spatial differences in infection will be more pronounced in *K. limnotrissae* as it infects only *L. miodon*;
2. there is differential occurrence of both species of *Kapentagyrus* on *L. miodon* driven by host lifestyle. We base this hypothesis on previously published results on differences in introduction success of members of *Kapentagyrus* between different life stages of these clupeid hosts (Kmentová et al., 2019);
3. given the contrasting patterns previously observed in cases of co-infection (Šimková et al. 2000, Hellard et al. 2015, Gobbin et al. 2021), both facilitation and competition on resources may be hypothesised between the species of *Kapentagyrus* infecting *L. miodon.* An overview and argumentation of the hypotheses is presented in Table 1.

**Table 1:**
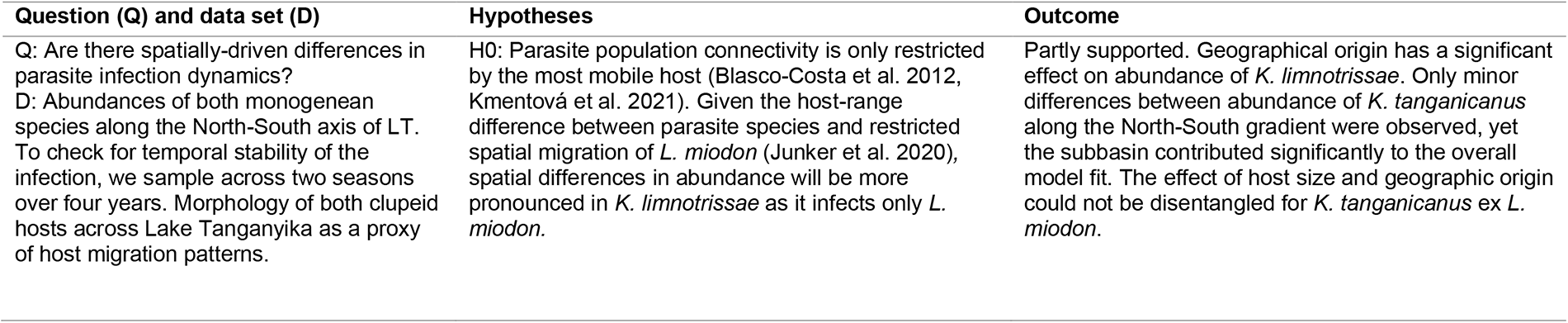
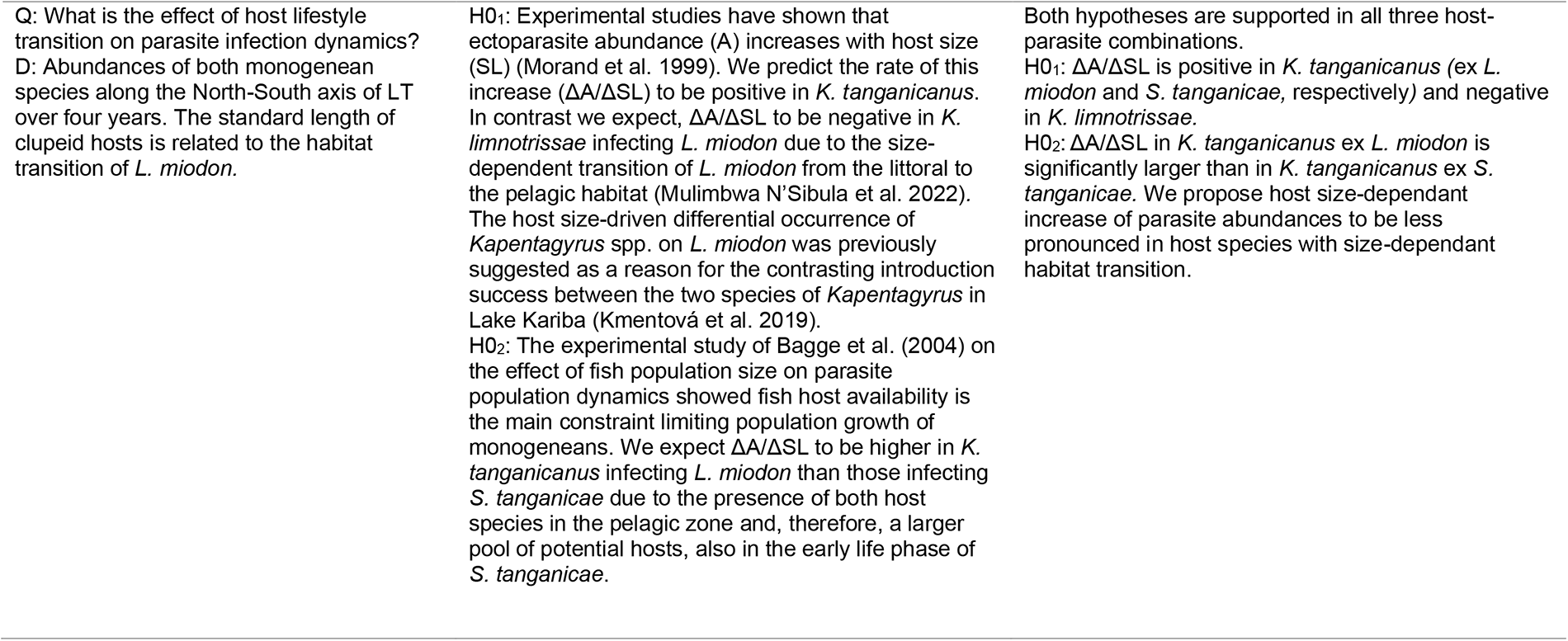
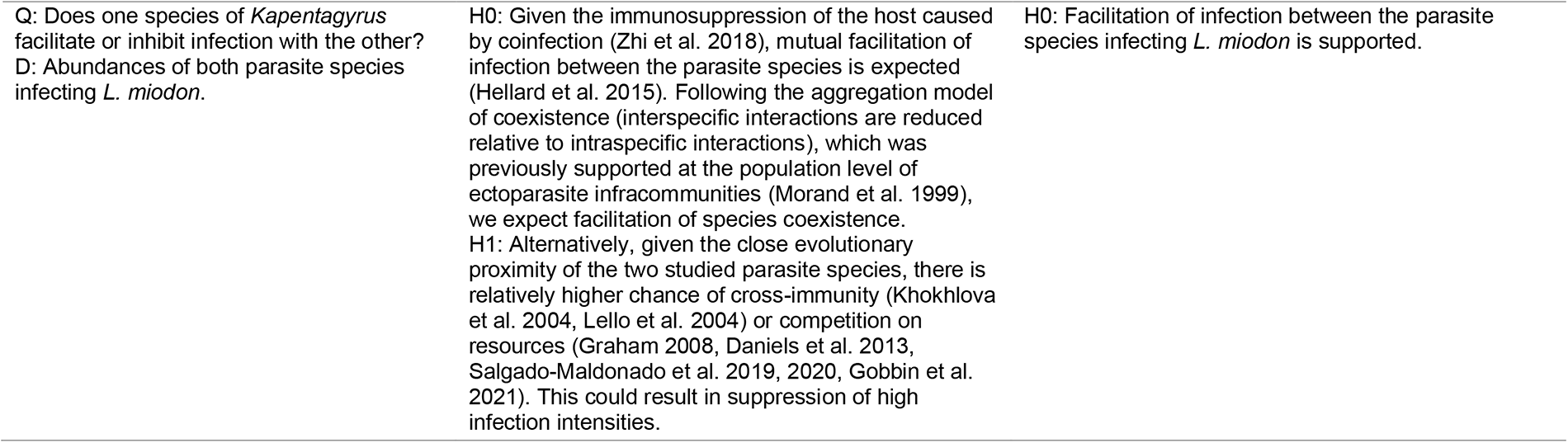
Overview of analysed research questions (Q), data sets (D), and hypotheses (H) with argumentation and outcomes for the studied host-parasite system (Fig 1).

## Methodology

### Sample collection and species identification

In total, 1730 specimens of pelagic clupeids, *L. miodon* (733) and *S. tanganicae* (997), were collected along the North-South axis of LT, including three subbasins (North, Central, and South) in two different seasons (rainy season from October to April and dry season from May to September) within a four-year period (Fig. 1, Table S1). Freshly caught fish specimens were either obtained with the experimental fishing unit of the Centre de Recherche en Hydrobiologie – Uvira (CRH) (Uvira, Democratic Republic of the Congo) or purchased from local fishermen. We combine newly obtained data on monogenean infection of clupeids in LT with those published in previous studies (Kmentová et al. 2016, 2018). Host specimens were collected within a period of two weeks (August 2016, April 2018, and October 2019) to avoid sampling the same population twice. This because both clupeid species are highly mobile (Mulimbwa N’Sibula and Mannini 1993, De Keyzer et al. 2019). Whole fish and/or gills were preserved in absolute ethanol. Host specimens were examined for the presence of monogenean parasites according to the procedure described in Kmentová et al. (2018). Species level identification of *Kapentagyrus* spp. was based on distinctive characters of the hard parts of the attachment organ in the posterior part of their bodies (for more details see Kmentová et al., 2018). In the case of *L. miodon*, the only species of the two clupeids that hosts two monogenean species, monogenean individuals that could not be identified at the species level because of damaged attachment organs (n=404) were only included in the counts of total infection intensities.

**Fig. 1:**
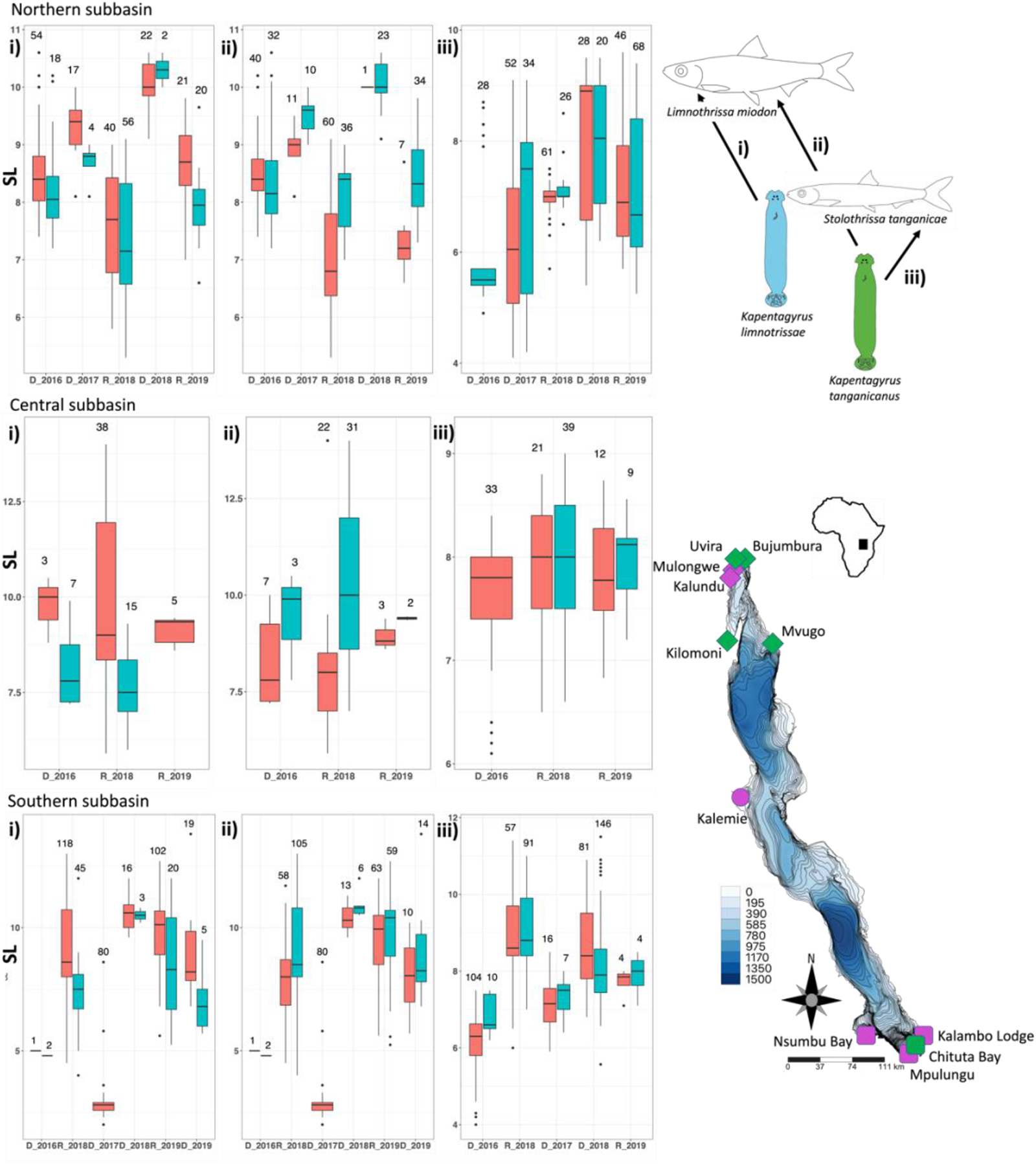
Sampling design and reported abundances of respective parasite populations. Season (Dry-D, Rainy – W) and year of origin (x-axis) and standard length of the fish host (y-axis, SL in cm) of i) *Kapentagyrus limnotrissae* ex *Limnothrissa miodon*, ii) *Kapentagyrus tanganicanus* ex *Limnothrissa miodon*, iii) *Kapentagyrus tanganicanus* ex *Stolothrissa tanganicae*. Infected fish are depicted in blue, non-infected in red. The total number of fish screened at a certain time point is mentioned above each of the boxplots. Subbasin division is indicated by different shapes used for sampling localities, with diamonds representing the northern subbasin, a circle representing the central subbasin and squares representing the southern subbasin. Sampling localities of fish specimens also used in the geomorphometric analyses are indicated in purple, those not used for geomorphometry in green. Bathymetry of the lake is presented in metres.

### Parasite population dynamics

To investigate the host parameters that might influence parasite abundance, the dataset was divided into three subsets based on host-parasite combinations according to host species (i: *K. limnotrissae* ex *L. miodon*, ii: *K. tanganicanus* ex *L. miodon*, iii: *K. tanganicanus* ex *S. tanganicae*) (see Fig. 1). We modelled the average infection intensity (i.e., per infected host specimen) against a range of other parameters, including sampling location (as subbasin – North, Central, South), season (dry period, rainy period), and host size (standard length) as explanatory variables (Table 2) as well as infection levels of the respective other parasite species if applicable, i.e. *K. limnotrissae* (i) and *K. tanganicanus* (ii) ex *L. miodon*. Because of seasonal migration and previous records on spatiotemporal variation in body size (Plisnier et al. 2009), we expected that host size would interact with locality and season. Several studies also suggested that infection levels of monogeneans are related to fish size (Akoll et al., 2012; Šimková et al., 2004). Therefore, we included host size interaction effects with the remaining parameters (locality, season) in the initial models.

Infection parameters of parasites often present a substantial amount of zero counts (Lester 2012, Tinsley et al. 2020). Therefore, we fitted infection levels using zero-inflated models (ZIMs) with a Poisson probability distribution, which assumes that the excess of zero counts is produced by a separate process. In the present case, we hypothesised that the excess of zero counts results from a lack of contact with parasites in some populations, while true zeros arise from host resistance (Wang, et al., 2017; Zuur et al., 2009). We fitted generalised linear models without zero-inflation to test whether these assumptions are true. The present datasets are overdispersed (residual deviance/residual degrees-of-freedom > 1.5 for a Poisson distribution). To address this overdispersal, we used a negative binomial probability distribution. Finally, host specimens most likely represent non-independent samples, as fish belonging to the same schools may have experienced similar parasite-exposure scenarios. Therefore, we also tested whether including the sampling day and locality as random effects in a mixed model further improved the model fit. To avoid overfitting, we simplified models through a backwards selection procedure using the function *drop1* including a χ² test. All effects that failed to significantly improve model fit were removed, starting from the interaction effects.

All model-based analyses were carried out in R v4.1.2 (R Core Team, 2022). Models were fitted using *glmmTMB* v1.1.2.3 (Brooks et al. 2017). This package offers two options for negative binomial distributions (options *nbinom1* and *nbinom2*) that implement linear and quadratic parameterisation, respectively (see Hardin & Hilbe, 2007). Both options were tested. We compared model fits using the Akaike information criterion (AIC) through the function *AICtab* in the package *bbmle* v1.0.24 (Bolker 2017). We also checked model fits through quantile-quantile plots and residual vs. fitted plots as provided by the package *DHARMa* v0.4.5 (Hartig 2022). We inferred changes of abundance over host standard length (ΔA/ΔSL) including the 95% confidence intervals from the model output in R. Intervals of ΔA/ΔSL that showed no overlap were considered significantly different from each other. Based on the best-fitting model, we predicted infection levels for all three host-parasite combinations as a function of the subbasin, as well as the continuous variables host standard length and the level of co-infections through the package *emmeans* (Lenth 2022). The resulting figures were plotted through the packages *emmeans* and *ggplot2* (Wickham 2016).

### Geomorphometrics of clupeid hosts

We verified partially restricted migration of clupeid hosts along the North-South axis of LT (De Keyzer et al. 2019, Junker et al. 2020) using geomorphometrics (Elewa 2004, Moreira et al. 2020). Migration patterns of clupeids could further influence infection dynamics of the two studied parasite species along the North-South axis of the lake. In October 2019, specimens were collected from six different localities along the lake’s shoreline within a period of two weeks (Fig. 1 and Table S1). Specimens were photographed using a Canon 4000D reflex camera equipped with an EF-S 18-55 mm III-lens, set on 55 mm for a total of 224 specimens of *S. tanganicae* and 195 specimens of *L. miodon*. The head shape, which was shown to be informative in studying phenotypic plasticity and population structure of clupeid fishes (Višnjić-Jeftić et al. 2013, Macossay-Cortez et al. 2022), was captured by a set of 12 fixed landmarks on each specimen. Landmarks were set using the *tpsDig2* software v2.31 (Rohlf, 2018) using a tps file created with the *tpsUtil* software v1.78 (Rohlf, 2018). The landmarks were defined based on previous studies conducted with other species of clupeids (Silva 2003, De La Cruz Agüero and Rodríguez 2004, Mounir et al. 2019), see Fig. S1. Morphological variation within the two clupeid species was analysed with *MorphoJ* v2 (Klingenberg 2011) (for more details see Supplementary File 1). Canonical Variate Analyses (CVA) on the residuals (see Supplementary File 1) and permutation tests of 10,000 replicates, were performed to test for differences in morphology between specimens from different sites of origin or subbasins. The resulting figures were plotted using the *R* packages *ggplot2*, *RColorBrewer* (Neuwirt 2022)*, ggtext* (Wilke 2020) and *tidyverse* (Wickham et al.

2019).

## Results

### Parasite population dynamics

For all three combinations (i–iii), a zero-inflated negative binomial mixed model had the best fit (Table 2). Seasonality failed to significantly improve model fit in all cases and this parameter was hence omitted, along with its interaction effects. The three minimal adequate models can be found in Table 2.

For *K. limnotrissae* (i), the models with a linear parameterisation (*nbinom1*) outperformed the quadratic parameterisation. The interaction effect of subbasin with standard length did not improve the model fit and was, therefore, removed. Abundances in the South of LT were significantly lower than in other subbasins (Fig. 2). Infections significantly decreased with host standard length and increased with co-infection intensities of *K. tanganicanus* (Fig. 2). For *K. tanganicanus* (ii and iii), a monogenean infecting both species of clupeids, the models with a quadratic parameterisation (*nbinom2*) outperformed the models with a linear parameterisation. In the post-hoc analysis, we found only minor differences between infection levels of *K. tanganicanus* along the North-South axis (Figs. 3&4). However, subbasin identity contributed significantly to the overall model fit in *K. tanganicanus* ex *L. miodon* (ii) (Table 2). Abundances of *K. tanganicanus* increased with host size of both clupeid species (Figs. 3&4) and with co-infection intensities of *K. limnotrissae* (Fig. 3). For *K. tanganicanus* ex *L. miodon* (ii), the increase with host size was generally weaker in the southern subbasin and particularly strong in the North. Host standard length had a significant interaction with the subbasin (Fig. 3). For *K. tanganicanus* ex *S. tanganicae* (iii), the standard lengths of the hosts were the only determinants of infection intensity (Fig. 4). In the minimum adequate models, ΔA/ΔSL is negative for *K. limnotrissae* (i) and positive for both host-parasite combinations of *K. tanganicanus* (ii, iii), yet ΔA/ΔSL for *K. tanganicanus* ex. *L. miodon* (ii) in the Central (0.61, CI: 0.43, 0.79) and Northern (0.86, CI: 0.33, 1.39) subbasins are significantly larger than for *K. tanganicanus* ex. *S. tanganicae* (0.16, CI: 0.04, 0.29).

**Table 2:**
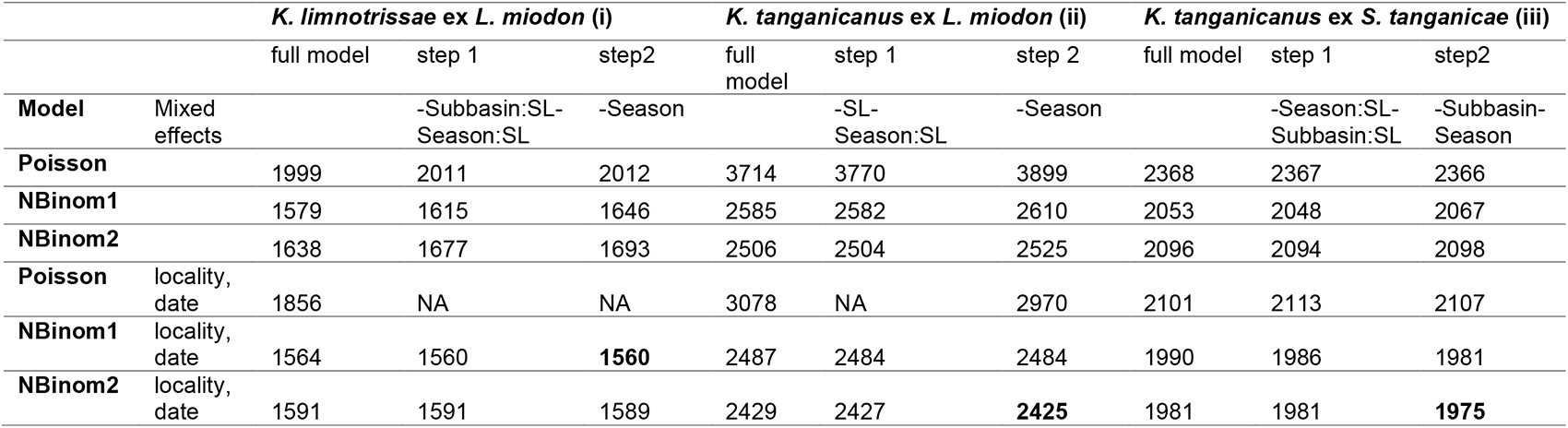
Stepwise backwards selection of effects (step 1 and 2) in generalised linear (mixed) models for each host-parasite combination. Models with the lowest AIC values (minimal adequate models) are highlighted in bold. Interaction terms are indicated with colons, removal of effects in indicated by minus sign.

|  |  | *K. limnotrissae* ex *L. miodon* (i) |  |  | *K. tanganicus* ex *L. miodon* (ii) |  |  | *K. tanganicus* ex *S. tanganicae* (iii) |  |  |
| --- | --- | --- | --- | --- | --- | --- | --- | --- | --- | --- |
|  |  | full model | step 1 | step2 | full model | step 1 | step 2 | full model | step 1 | step2 |
| **Model** | Mixed effects |  | -Subbasin:SL-Season:SL | -Season |  | -SL-Season:SL | -Season |  | -Season:SL-Subbasin:SL | -Subbasin-Season |
| **Poisson** |  | 1999 | 2011 | 2012 | 3714 | 3770 | 3899 | 2368 | 2367 | 2366 |
| **NBinom1** |  | 1579 | 1615 | 1646 | 2585 | 2582 | 2610 | 2053 | 2048 | 2067 |
| **NBinom2** |  | 1638 | 1677 | 1693 | 2506 | 2504 | 2525 | 2096 | 2094 | 2098 |
| **Poisson** | locality,<br>date | 1856 | NA | NA | 3078 | NA | 2970 | 2101 | 2113 | 2107 |
| **NBinom1** | locality,<br>date | 1564 | 1560 | **1560** | 2487 | 2484 | 2484 | 1990 | 1986 | 1981 |
| **NBinom2** | locality,<br>date | 1591 | 1591 | 1589 | 2429 | 2427 | **2425** | 1981 | 1981 | **1975** |

**Fig. 2:**
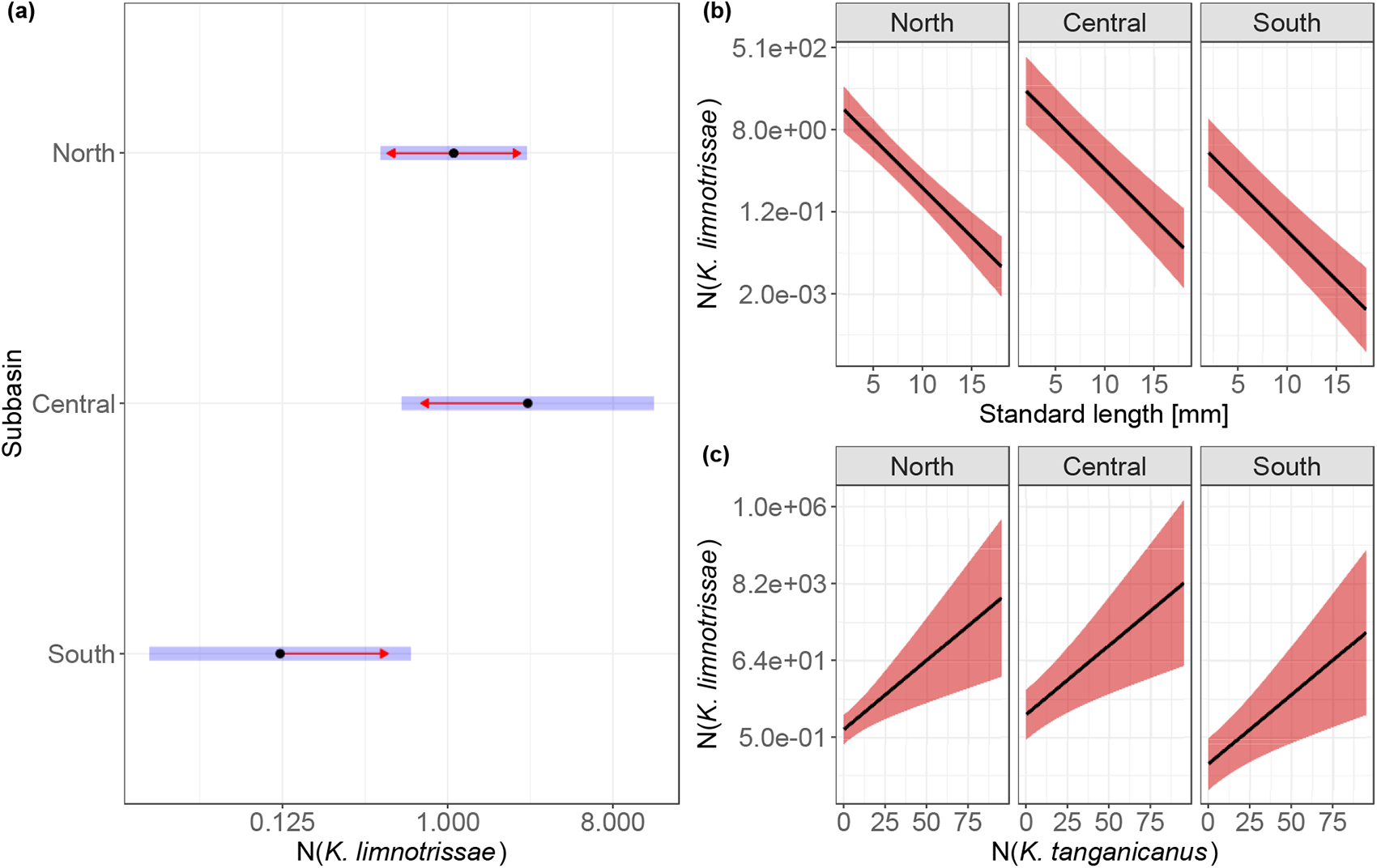
Predicted abundance of *Kapentagyrus limnotrissae* ex *Limnothrissa miodon*. a) Infection intensity related to subbasin origin with the confidence interval (in blue) and direction of overlap (in red), b) Infection intensity as a function of the standard length (SL) of the host, c) Infection intensity as function of co-infection by *Kapentagyrus tanganicanus* (infection intensity on x-axis).

**Fig. 3:**
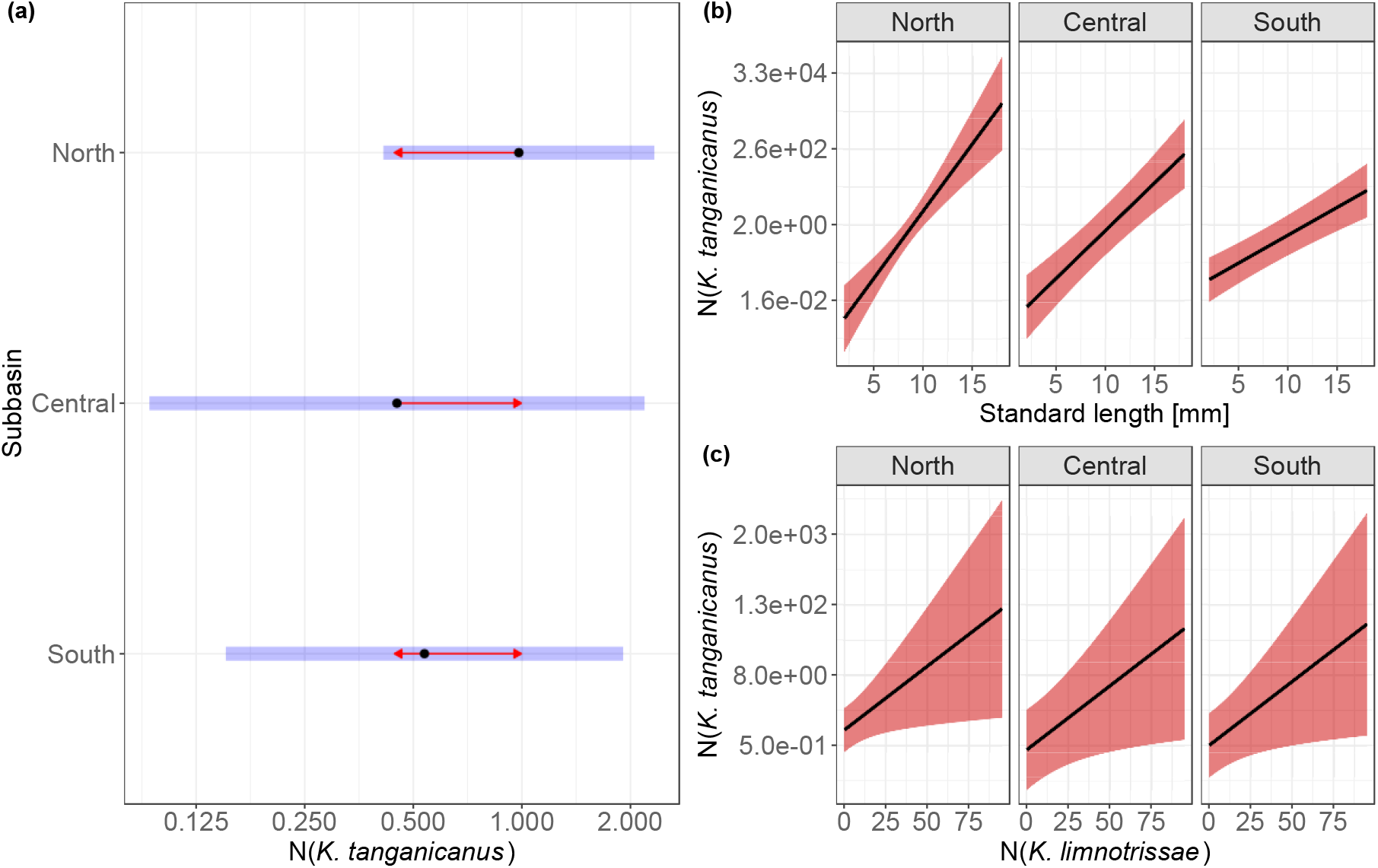
Predicted abundance of *Kapentagyrus tanganicanus* ex *Limnothrissa miodon* with 95% confidence intervals under the minimal adequate model (Table 2). a) Infection intensity in different subbasins showed no significant differences (CIs in blue, direction of overlap in red), b) infection intensity decreased with standard length (SL) of the host, c) infection intensity increased with higher infection intensities of *Kapentagyrus limnotrissae*.

**Fig. 4:**
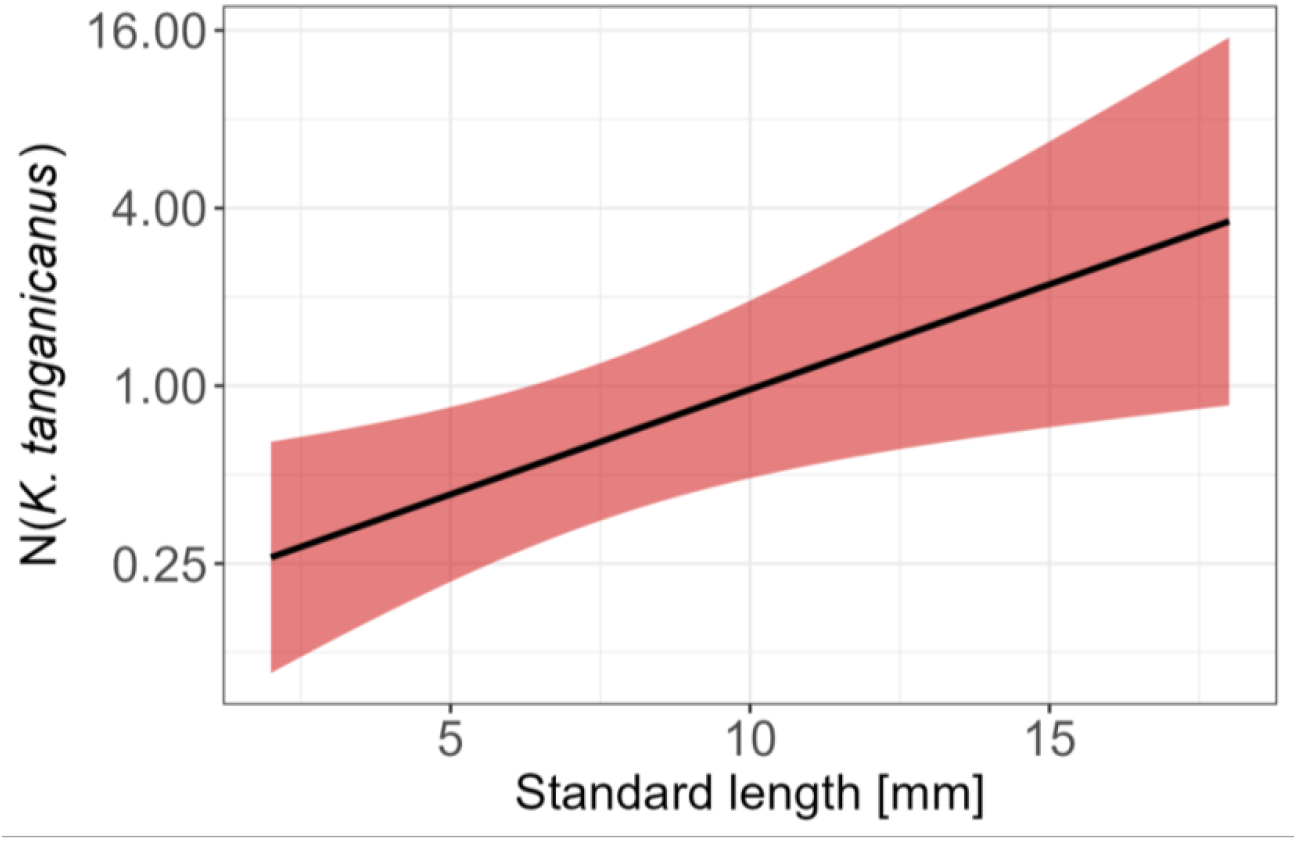
Predicted abundance of *Kapentagyrus tanganicanus* ex *Stolothrissa tanganicae*. Infection intensity increased with the standard length (SL) of the host.

### Geomorphometrics of clupeid hosts

For both species of clupeids, geomorphometric analyses revealed a strong similarity in head morphology between specimens from different localities and subbasins of the lake. In the case of *L. miodon*, the first two PC axes explained 22.1% and 17.1% of the variation, respectively (see Fig. 5a&b). In case of *S. tanganicae*, the first two PC axes explained 24.2% and 18.1% of the variation, respectively (see Fig. 5c&d). Based on the wireframes (Fig. S2), differences in the relative position of the snout, eye and operculum are visible along the displayed PC axes.

**Fig. 5:**
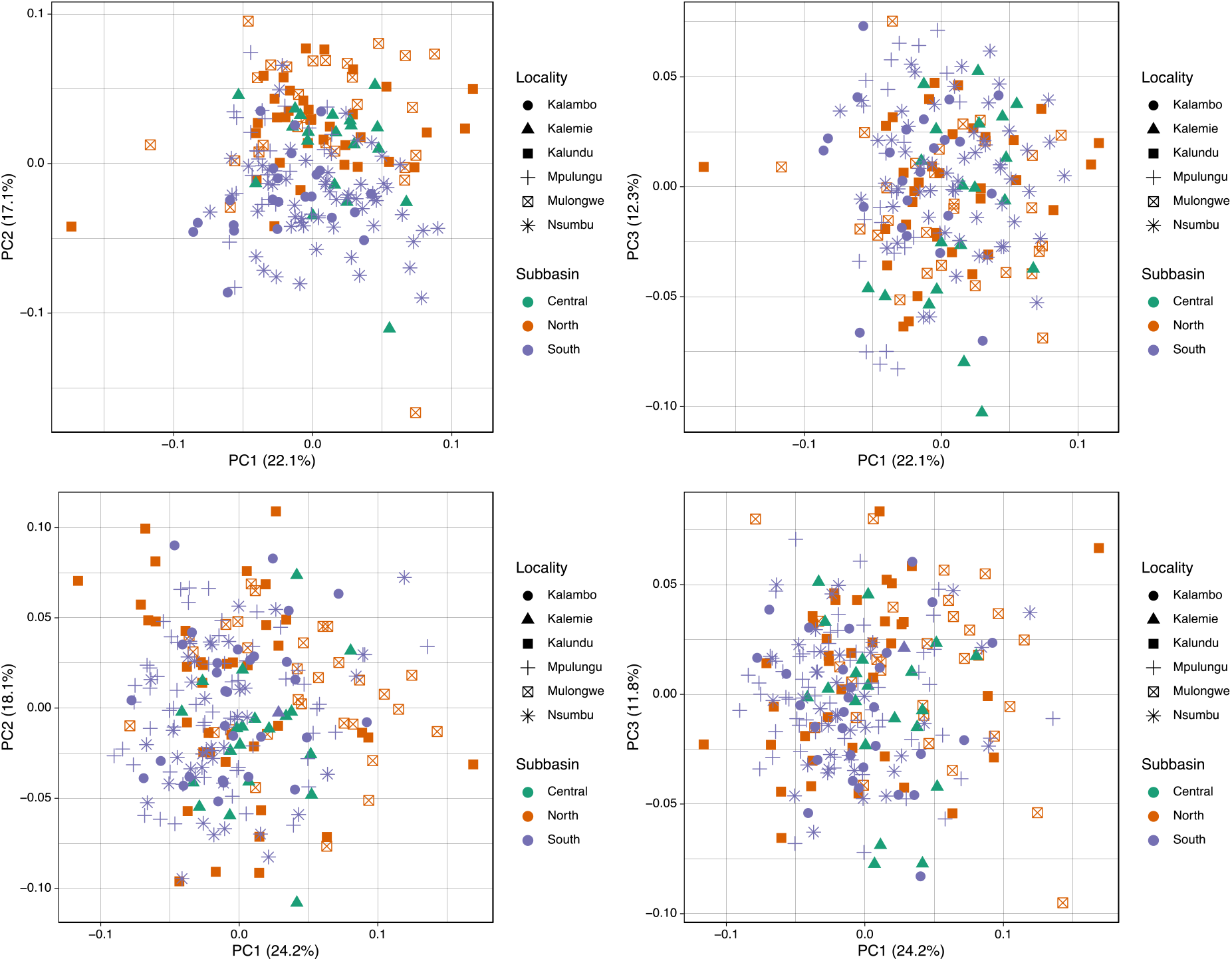
Biplots of Principal Component Analyses (PCA) showing the shape variation in the head across the sampled localities of a) *Limnothrissa miodon*, first two PCs displayed, b) *Limnothrissa miodon*, first and third PCs displayed, c) *Stolothrissa tanganicae,* first two PCs displayed, d) *Stolothrissa tanganicae,* first and third PCs displayed.

The results of our CVAs show significant differences in the head shape related to the geographic origin of *L. miodon* (Table 3), a trend visible mainly along the second PC axis (Figure 6). Although the CVAs indicated significant differences in shape between some of the localities of *S. tanganicae* (Table 3), no consistent geographical pattern was detected in the PCAs.

**Table 3:** Results of Canonical Variate Analyses of head shape variation for a) *Limnothrissa miodon* b) *Stolothrissa tanganicae*. Values of Procrustes distances are displayed below the diagonal, P-values are shown above the diagonal. Significant P-values (< 0.05) are indicated in bold. Numbers of fish specimens are indicated between brackets behind the locality names (column names), subbasin is specified behind the locality (row names): N – North, C – Central, S – South.

| **(a)** | Mulongwe (28) | Kalundu (34) | Kalemie (20) | Mpulungu (27) | Nsumbu (58) | Kalambo (23) |
| --- | --- | --- | --- | --- | --- | --- |
| Mulongwe (N) | x | 0.3842 | **0.0207** | **0.0002** | **0.0466** | **0.0001** |
| Kalundu (N) | 0.0252 | x | **0.0029** | **0.0002** | **0.0376** | **0.0001** |
| Kalemie (C) | 0.0447 | 0.0423 | x | **<.0001** | **0.0355** | **0.0005** |
| Mpulungu (S) | 0.0519 | 0.0472 | 0.0550 | x | **0.0027** | 0.2593 |
| Nsumbu (S) | 0.0298 | 0.0263 | 0.0308 | 0.0340 | x | **0.0008** |
| Kalambo (S) | 0.0603 | 0.0536 | 0.0489 | 0.0264 | 0.0379 | x |
| **(b)** | Mulongwe (30) | Kalundu (38) | Kalemie (21) | Mpulungu (64) | Nsumbu (40) | Kalambo (30) |
| Mulongwe (N) | x | **0.0060** | 0.0570 | **0.0029** | **<.0001** | **0.0014** |
| Kalundu (N) | 0.0479 | x | 0.0651 | 0.5339 | 0.0321 | 0.1954 |
| Kalemie (C) | 0.0387 | 0.0401 | x | **0.0191** | **0.0116** | **0.0336** |
| Mpulungu (S) | 0.0382 | 0.0188 | 0.0336 | x | **0.0196** | 0.3979 |
| Nsumbu (S) | 0.0584 | 0.0339 | 0.0368 | 0.0270 | x | 0.0785 |
| Kalambo (S) | 0.0467 | 0.0301 | 0.0353 | 0.0189 | 0.0274 | x |

## Discussion

### Spatial dynamics of parasite infection – effect of the most mobile host

We found a difference in spatial structure between the three host-parasite combinations, confirming that distributions of *Kapentagyrus* spp. are shaped by the migration patterns of their respective most mobile host species. Previous studies indicate that the distribution of aquatic parasites is determined by environmental factors (Timi and Poulin 2003). Here, we found no effect of seasonality on the spatiotemporal dynamics of *Kapentagyrus* spp. This in spite of the reported North-South clupeid migrations, resulting in seasonal fluctuations of host population densities and mean size distribution (Kimirei and Mgaya 2007, Mulimbwa N’Sibula et al. 2022). This result confirms the previously suggested independence of seasonal host population cycles of rather short-lived parasites (Lester and MacKenzie 2009, Henriksen et al. 2023), among which monogeneans in tropical areas may be counted with their estimated generation time of days up to a few weeks (Tomnatik 1990). In contrast to our study system, host behaviour was shown to drive temporal differences in infection levels of *Mazocraes alosae* on the clupeid *Alosa immaculata* in the northern Black Sea and the Sea of Azov (Plaksina et al. 2021).

While we could not fully disentangle the effects of host size and locality for *K. tanganicanus* ex *L. miodon*, overlapping abundances were observed in all three subbasins. *Kapentagyrus limnotrissae* exhibited a slight spatial differentiation in infection along the North-South axis. Previously, restricted migration of *L. miodon* compared to *S. tanganicae* was evidenced by Phiri and Shirakihara (1999) and Mulimbwa N’Sibula et al. (2014) and recent genomic patterns along the North-South axis of LT (De Keyzer et al. 2019, Junker et al. 2020). This is confirmed by our geomorphometric analyses, indicating a more profound head shape differentiation in *L. miodon* compared to *S. tanganicae,* even while originating from the same localities. We suggest that this pattern is driven by temporal residency and patchy distribution of *L. miodon* in the littoral habitat before reaching a certain size and becoming pelagic. Together, the lack of spatial differences of infection in *K. tanganicanus* and the correspondence of restricted spatial distribution in *K. limnotrissae* and *L. miodon* support the general ecological principle that parasite distribution is restricted by the most mobile host (Blasco-Costa et al. 2012, Kmentová et al. 2021). In the pelagic zone of LT, similar result was also reported for *Cichlidogyrus casuarinus* infecting seven cichlid species (Kmentová et al., 2021).

In comparison to monogeneans infecting clupeid fishes in LT, limited spatial distribution of the monogenean parasite compared to the host of *Pseudanthocotyloides heterocotyle* infecting *Sprattus sprattus* (Kleinertz et al., 2012) and *Clupea harengus* (Actinopterygii, Clupeidae) (Rahimian et al., 1999) in the North Sea was observed.

### Infection dynamics driven by host size and lifestyle

The body size of the clupeid hosts appears to be a major determinant of monogenean infection in LT, as it significantly affects abundances for all three host-parasite combinations. We found a positive correlation between parasite abundance and host size in two (ii, iii) and a negative correlation in one (i) of the three host-parasite combinations (Table 1). A positive correlation between fish size and monogenean abundance has been associated with the larger habitat offered by larger hosts (Alvarez-Pellitero & Gonzalez-Lanza, 1982; Poulin, 2000). Considering the age-dependent migration of *L. miodon* from the littoral to the pelagic zone, the contrasting pattern of occurrence between *K. limnotrissae* ex *L. miodon* (i) and *K. tanganicanus* ex *L. miodon* (ii) associated with host size (reflecting age differences) suggests a spatial stratification of infection linked with the ontogenetic migration of *L. miodon* (Coulter 1970, 1991b, Mannini et al. 1996, Mulimbwa N’Sibula et al. 2022). This corresponds to the previous reports of clupeid introductions in Lake Kariba, where differential co-introduction success of *Kapentagyrus* spp. on *L. miodon* was observed and related to limited size of translocated specimens of the host (Kmentová et al., 2019).

The results of an experimental study by Bagge et al. (2004) support the positive relationship between the fish population size and parasite infection. An overall higher host species richness and population size of clupeids in the pelagic zone (in contrast to the littoral zone occupied by smaller-sized *L. miodon* only) could further explain the smaller rate of increase in abundances of *K. tanganicanus* on *S. tanganicae* (iii) compared to on *L. miodon* (ii) in the pelagic zone. To our knowledge this is the first time the effect of host species richness and/or population size on fish parasite population dynamics is proposed in natural conditions.

### Facilitation of infection – sharing is beneficial

The apparent host-size related replacement in the two parasite species infecting *L. miodon* (i, ii) contrasts with the beneficial influence that each of the parasite species has on the other (Table 1, Figs 2&3). Interactions among parasites infecting the same host individual are greatly understudied in natural populations. In the literature, the mode of transmission, virulence and possible host manipulation are listed as important factors driving co-infection dynamics (Haine et al. 2005, Rigaud et al. 2010). Evolutionarily closely-related parasites with the same mode of transmission are more likely to benefit from each other’s influence on their host. This was reported for microsporidian and paramyxean parasites infecting an amphipod host (Short et al. 2012, Arundell et al. 2015, Pickup and Ironside 2018). Previous studies on monogeneans revealed that suppressing the fish immune system allows for increased infection intensities across monogenean species (Sitjà-Bobadilla 2008, Rohlenová et al. 2011) as well as other parasite taxa (Klemme et al., 2016). This effect could explain facilitation of infection between the species of *Kapentagyrus* on *L. miodon*. So far, no evidence of positive or any direct interspecific interaction between closely related monogenean species has been found (Šimková et al., 2000; Soler-Jiménez & Fajer-Ávila, 2012). This study is the first to show such a positive interaction in a natural environment.

Additionally, it is the first to show it co-occurring with a lifestyle-dependent replacement of one parasite species by another.

## Conclusions

Only a few general patterns in the ecology of macroparasites from natural ecosystems have been established. The lack of confounding factors in the ecologically, geographically and evolutionarily well-delineated clupeid-monogenean host-parasite system in LT makes it ideal to study large scale patterns of parasites distribution.

Differences in migration patterns between the fish host species are suggested to drive spatial population dynamics of monogenean parasites on a lake-wide scale. Our results therefore support general dependence of parasite population dynamics on spatial distribution of their most mobile host species. Contrasting occurrence patterns of the two parasite species on a shared fish host species further illustrate the dependence on changes in lifestyle of the host during its ontogenetic development and support the previously suggested importance of host body size for the parasites’ introduction success in non-native areas. The evidenced facilitation of infection between the two studied parasite species further spins the discussion on the direction of parasite-parasite interactions.

## Acknowledgements

We would like to thank Stephan Koblmüller, Holger Zimmermann, Jiří Vorel, Simona Georgieva, Gyrhaiss Kapepula Kasembele, Cyprian Katongo, Taylor Banda, Aneesh P.H. Bose, Filip A.M. Volckaert and Els De Keyzer for their help in organising and conducting field work. Pierre-Denis Plisnier is acknowledged for providing bathymetric data of Lake Tanganyika. The study was supported by the Czech Science Foundation (GACR) project no. P505/12/G112—European Centre of Ichtyoparasitology (ECIP) and standard project GA19-13573S, research grant 1513419N of the Research Foundation—Flanders (FWO-Vlaanderen), Global Minds project GM2O18INITO7 of Hasselt University, and South Initiative CD2018SIN218A101 of VLIR-UOS Travel grant (2019). AJCL (BOF19OWB02), NK (BOF21PD01) and MPMV (BOF20TT06) received support from the Special Research Fund of Hasselt University.

## Conflict of Interest

Authors declare no conflict of interest.

## Author Contributions

N.K. designed the study, generated incidence data, analysed geometric morphometric data and drafted the manuscript. A.C-L. analysed incidence data and helped draft the manuscript, M.W.P.J. generated part of the incidence data and helped to draft the manuscript, M.V.S. supervised geometric morphometric data analyses and interpretation of results, M.P.M.V. discussed the results, helped draft the manuscript and supervised the study, T.M.and E.V.H. contributed to geometric morphometric data analyses, S. H. helped with geometric morphometric data acquisition, L.Mi. and K.J.M.T. helped with data interpretation and drafting the manuscript, P.M.M., A.C.M. and L.Ma. provided support in the field and knowledge on the studied ecosystem. All authors have read and agreed to the published version of the manuscript.

## Data Availability Statement

Parasite voucher material was deposited in the collection of Hasselt University under accession numbers xx-xx and the Royal Museum for Central Africa under accession numbers xx-xx. The infection data and geometric morphometric data underlying the results of this article are available in Mendeley Data (doi: 10.17632/ntwrrtknyf.1).

## Supporting Information

Additional supporting information may be found in the online version of the article at the publisher’s website.

## Notes

https://data.mendeley.com/drafts/ntwrrtknyf

### Competing Interest Statement

The authors have declared no competing interest.

### Summary of Updates

Link questions with the concrete hypotheses.

## References

Akoll, P. et al. 2012. Risk assessment of parasitic helminths on cultured Nile tilapia (Oreochromis niloticus, L.). – Aquaculture 356–357: 123–127.

Alvarez-Pellitero, M. P. and Gonzalez-Lanza, M. C. 1982. Description and population dynamics of *Dactylogyrus legionensis* n. sp. from *Barbus barbus bocagei* Steind. – J. Helminthol. 56: 263–273.

Arundell, K. et al. 2015. Enemy release and genetic founder effects in invasive killer shrimp populations of Great Britain. – Biol. Invasions 17: 1439–1451.

Bagge, A. M. et al. 2004. Fish population size, and not density, as the determining factor of parasite infection: a case study. – Parasitology 128: 305–313.

Blackburn, T. M. et al. 2011. A proposed unified framework for biological invasions. – Trends Ecol. Evol. 26: 333–339.

Blasco-Costa, I. et al. 2012. Swimming against the current: genetic structure, host mobility and the drift paradox in trematode parasites. – Mol. Ecol. 21: 207–217.

Bolker, B. 2017. Package ‘bbmle’.“ Tools for General Maximum Likelihood Estimation.

Brooks, M. E.;, et al. 2017. glmmTMB balances speed and flexibility among packages for zero-inflated generalized linear mixed modeling. – R J. 9: 378–400.

Buck, J. C. and Lutterschmidt, W. I. 2017. Parasite abundance decreases with host density: evidence of the encounter-dilution effect for a parasite with a complex life cycle. – Hydrobiologia 784: 201–210.

Cote, I. M. and Poulin, R. 1995. Parasitism and group size in social animals: a meta-analysis. – Behav. Ecol. 6: 159–165.

Coulter, G. W. 1970. Population changes within a group of fish species in Lake Tanganyika following their exploitation. – J. Fish Biol. 2: 329–353.

Coulter, G. W. 1991a. Introduction. – In: Coulter, G. W. (ed), Lake Tanganyika and Its Life. Natural History Museum Publications & Oxford University Press, pp. 1–6.

Coulter, G. W. 1991b. Pelagic Fish. – In: Coulter, G. W. (ed), Lake Tanganyika and Its Life. Natural History Musem & Oxford University Press, pp. 111–150.

Criscione, C. D. and Blouin, M. S. 2004. Life cycles shape parasite evolution: comparative population genetics of salmon trematodes. – Evolution (N. Y). 58: 198–202.

Cristescu, M. E. et al. 2010. Ancient lakes revisited: from the ecology to the genetics of speciation. – Mol. Ecol. 19: 4837–4851.

Daniels, R. R. et al. 2013. Do parasites adopt different strategies in different intermediate hosts? Host size, not host species, influences Coitocaecum parvum (Trematoda) life history strategy, size and egg production. – Parasitology 140: 275–283.

De Keyzer, E. L. R. et al. 2019. First genomic study on Lake Tanganyika sprat *Stolothrissa tanganicae*: A lack of population structure calls for integrated management of this important fisheries target species. – BMC Evol. Biol. 19: 6.

De La Cruz Agüero, J. and Rodríguez, F. J. G. 2004. Morphometric stock structure of the Pacific sardine Sardinops sagax (Jenyns, 1842) off Baja California, Mexico. – In: Elewa, A. M. T. (ed), Morphometrics: Applications in Biology and Paleontology. Springer Berlin Heidelberg, pp. 115–127.

El Hafidi, F., et al. 1998. Microhabitat distribution and coexistence of Microcotylidae (Monogenea) on the gills of the striped mullet *Mugil cephalus*: Chance or competition? – Parasitol. Res. 84: 315–320.

Elewa, A. 2004.Morphometrics: applications in biology and paleontology. Vol.14. – Springer Science & Business Media.

Gobbin, T. P. et al. 2021. Microhabitat distributions and species interactions of ectoparasites on the gills of cichlid fish in Lake Victoria, Tanzania. - Int. J. Parasitol. 51: 201–214.

Graham, A. L. 2008. Ecological rules governing helminth-microparasite coinfection. – Proc. Natl. Acad. Sci. U. S. A. 105: 566–570.

Haine, E. R. et al. 2005. Conflict between parasites with different transmission strategies infecting an amphipod host. – Proc. R. Soc. B Biol. Sci. 272: 2505–2510.

Hardin, J. W. and Hilbe, J. M. 2007. Generalized linear models and extensions. – Stata press.

Hartig, F. 2022. DHARMa: Residual Diagnostics for Hierarchical (Multi-Level / Mixed) Regression Models. R package version 0.4.6.

Hatcher, M. J. et al. 2012. Disease emergence and invasions. – Funct. Ecol. 26: 1275–1287.

Hellard, E. et al. 2015. Parasite-parasite interactions in the wild: how to detect them? – Trends Parasitol. 31: 640–652.

Henriksen, E. H. et al. 2023. Ectoparasites population dynamics are affected by host body size but not host density or water temperature in a 32-year long time series. – Oikos 2023: 1–13.

Junker, J. et al. 2020. Structural genomic variation leads to genetic differentiation in Lake Tanganyika’s sardines. – Mol. Ecol. 29: 3277–3298.

Kennedy, C. R. 2001.Interspecific interactions between larval digeneans in the eyes of perch, *Perca fluviatilis*. – Parasitology 122: S13–S22.

Khokhlova, I. S. et al. 2004. Immune response to fleas in a wild desert rodent: effect of parasite species, parasite burden, sex of host and host parasitological experience. – J. Exp. Biol. 207: 2725–2733.

Kimirei, I. and Mgaya, Y. 2007. Influence of environmental factors on seasonal changes in clupeid catches in the Kigoma area of Lake Tanganyika. – African J. Aquat. Sci. 32: 291–298.

Kleinertz, S. et al. 2012. Parasite communities and feeding ecology of the European sprat (*Sprattus sprattus* L.) over its range of distribution. – Parasitol. Res. 110: 1147–1157.

Klemme, I. et al. 2016. Host infection history modifies co-infection success of multiple parasite genotypes. – J. Anim. Ecol. 85: 591–597.

Klingenberg, C. P. 2011. MorphoJ: an integrated software package for geometric morphometrics. – Mol. Ecol. Resour. 11: 353–357.

Kmentová, N. 2019. The parasite fauna of economically important pelagic fishes in Lake Tanganyika. Doctoral thesis, Masaryk University, Brno, Czech Republic.

Kmentová, N. et al. 2016. Reduced host-specificity in a parasite infecting non-littoral Lake Tanganyika cichlids evidenced by intraspecific morphological and genetic diversity. – Sci. Rep. 6: 39605.

Kmentová, N. et al. 2018. Monogenean parasites of sardines in Lake Tanganyika: diversity, origin and intra-specific variability. – Contrib. to Zool. 87: 105–132.

Kmentová, N. et al. 2019. Co-introduction success of monogeneans infecting the fisheries target *Limnothrissa miodon* differs between two non-native areas: The potential of parasites as a tag for introduction pathway. – Biol. Invasions 21: 757–773.

Kmentová, N. et al. 2021. Contrasting host-parasite population structure: Morphology and mitogenomics of a parasitic flatworm on pelagic deepwater cichlid fishes from Lake Tanganyika. – Biology (Basel). 10: 797.

Kmentová, N. et al. 2022. Dactylogyridae 2022: A meta-analysis of phylogenetic studies and generic diagnoses of parasitic flatworms using published genetic and morphological data. – Int. J. Parasitol. 52: 427–457.

Kuris, A. M. et al. 2008. Ecosystem energetic implications of parasite and free-living biomass in three estuaries. – Nature 454: 515–518.

Lafferty, K. D. et al. 2006. Parasites dominate food web links. – Proc. Natl. Acad. Sci. 103: 11211–11216.

Lefèvre, T. et al. 2009. The ecological significance of manipulative parasites. – Trends Ecol. Evol. 24: 41–48.

Lello, J. et al. 2004. Competition and mutualism among the gut helminths of a mammalian host. – Nature 428: 840–844.

Lenth, R. V. 2022. emmeans: Estimated marginal means (Least-squares means). R package version 1.7.5.

Lester, R. J. G. 2012. Overdispersion in marine fish parasites. – J. Parasitol. 98: 718–721.

Lester, R. J. G. and MacKenzie, K. 2009. The use and abuse of parasites as stock markers for fish. – Fish. Res. 97: 1–2.

Lima, D. P. et al. 2012. Patterns of interactions of a large fish–parasite network in a tropical floodplain. – J. Anim. Ecol. 81: 905–913.

Luque, J. L. et al. 2004. Parasite biodiversity and its determinants in coastal marine teleost fishes of Brazil. – Parasitology 128: 671–682.

Macossay-Cortez, A. et al. 2022. Intraspecific morphological variation in shads, Dorosoma anale and D. petenense (Actinopterygii: Clupeiformes: Clupeidae), in the Mexican Grijalva and Usumacinta river basins. – Acta Ichthyol. Piscat. 52: 149–158.

Mannini, P. et al. 1996. Pelagic fish stocks of Lake Tanganyika: Biology and exploitation. FAO/FINNIDA research for the management of the fisheries of Lake Tanganyika. – GCP/RAF/271/FIN—TD/53 (En)

Marcogliese, D. J. 2002. Food webs and the transmission of parasites to marine fish. – Parasitology 124: 83–99.

Milec, L. J. M. et al. 2022. Complete mitochondrial genomes and updated divergence time of the two freshwater clupeids endemic to Lake Tanganyika (Africa) suggest intralacustrine speciation. – BMC Ecol. Evol. 22: 1–17.

Morand, S. et al. 1999. Aggregation and species coexistence of ectoparasites of marine fishes. – Int. J. Parasitol. 29: 663–672.

Moreira, C. et al. 2020. Landmark-based geometric morphometrics analysis of body shape variation among populations of the blue jack mackerel, Trachurus picturatus, from the North-East Atlantic. – J. Sea Res. 163: 101926.

Mounir, A. et al. 2019. Discrimination of the phenotypic sardine *Sardina pilchardus* stocks off the Moroccan Atlantic coast using a morphometric analysis. – https://doi.org/10.2989/1814232X.2019.1597765 41: 137–144.

Mulimbwa N’Sibula, T. and Mannini, P. 1993. Demographic characteristics of Stolothrissa tanganicae, Limnothrissa miodon and Lates stappersii in the Northwestern (Zairean) waters of Lake Tanganyika. – CIFA Occas. Pap. in press.

Mulimbwa N’Sibula, T., et al. 2014. Seasonal changes in the pelagic catch of two clupeid zooplanktivores in relation to the abundance of copepod zooplankton in the northern end of Lake Tanganyika. – Aquat. Ecosyst. Heal. Manag. 17: 25–33.

Mulimbwa N’Sibula, T., et al. 2022. Spatial and seasonal variation in reproductive indices of the clupeids *Limnothrissa miodon* and *Stolothrissa tanganicae* in the Congolese waters of northern Lake Tanganyika. – Belgian J. Zool. 152: 13–31.

Neuwirt, E. 2022. RColorBrewer: ColorBrewer Palettes. R package version 1.1-3. in press.

Paugy, D. and Lévêque, C. 2017. Diets and food webs. – In: Paugy, D. et al. (eds), The inland water fishes of Africa: Diversity, ecology and human use. pp. 233–257.

Phiri, H. and Shirakihara, K. 1999. Distribution and seasonal movement of pelagic fish in southern Lake Tanganyika. – Fish. Res. 41: 63–71.

Pickup, J. and Ironside, J. E. 2018. Multiple origins of parasitic feminization: thelygeny and intersexuality in beach-hoppers are caused by paramyxid parasites, not microsporidia. – Parasitology 145: 408–415.

Plaksina, M. P. et al. 2021. Life-history studies on infrapopulations of *Mazocraes alosae* (Monogenea) parasitising *Alosa immaculata* (Actinopterygii) in the northern Black and Azov Seas. – http://folia.paru.cas.cz/doi/10.14411/fp.2021.009.html 68: 1–10.

Plisnier, P.-D. et al. 1999. Limnological annual cycle inferred from physical-chemical fluctuations at three stations of Lake Tanganyika. – In: Lindqvist, O. V. et al. (eds), From Limnology to Fisheries: Lake Tanganyika and Other Large Lakes. Springer Netherlands, pp. 45–58.

Plisnier, P. D. et al. 2009. Limnological variability and pelagic fish abundance (*Stolothrissa tanganicae* and *Lates stappersii*) in Lake Tanganyika. – Hydrobiologia 625: 117–134.

Poulin, R. 2000. Variation in the intraspecific relationship between fish length and intensity of parasitic infection: biological and statistical causes. – J. Fish Biol. 56: 123–137.

Poulin, R. 2003. The decay of similarity with geographical distance in parasite communities of vertebrate hosts. – J. Biogeogr. 30: 1609–1615.

Poulin, R. 2007. Are there general laws in parasite ecology? – Parasitology 134: 763–776.

Poulin, R. 2013. Explaining variability in parasite aggregation levels among host samples. - Parasitology 140: 541–546.

Poulin, R. 2014. Parasite biodiversity revisited: frontiers and constraints. – Int. J. Parasitol. 44: 581–589.

Poulin, R. 2020. Meta-analysis of seasonal dynamics of parasite infections in aquatic ecosystems. – Int. J. Parasitol. 50: 501–510.

Poulin, R. and Morand, S. 1999. Geographical distances and the similarity among parasite communities of conspecific host populations. – Parasitology 119: 369–374.

Rahimian, H. et al. 1999. Pseudanthocotyloides heterocotyle (van Beneden, 1871) Euzet & Prost, 1969 (Monogenea: Polyopisthocotylea: Mazocraeidae), a parasite of herring Clupea harengus L. and sprat Sprattus sprattus L. (Teleostei: Clupeidae). – Syst. Parasitol. 1999 423 42: 193–201.

Rigaud, T. et al. 2010. Parasite and host assemblages: embracing the reality will improve our knowledge of parasite transmission and virulence. – Proc. R. Soc. B Biol. Sci. 277: 3693–3702.

Rohlenová, K. et al. 2011. Are fish immune systems really affected by parasites? an immunoecological study of common carp (*Cyprinus carpio*). – Parasit. Vectors 4: 120.

Salgado-Maldonado, G. et al. 2019. Aggregation and negative interactions in low-diversity and unsaturated monogenean (Platyhelminthes) communities in *Astyanax aeneus* (Teleostei) populations in a neotropical river of Mexico. – Int. J. Parasitol. Parasites Wildl. 8: 203–215.

Salgado-Maldonado, G. et al. 2020. Competition from sea to mountain: Interactions and aggregation in low-diversity monogenean and endohelminth communities in twospot livebearer *Pseudoxiphophorus bimaculatus* (Teleostei: Poeciliidae) populations in a neotropical river. – Ecol. Evol. 10: 9115–9131.

Sasal, P. et al. 2008. Parasite communities in eels of the Island of Reunion (Indian Ocean): a lesson in parasite introduction. – Parasitol. Res. 102: 1343–1350.

Short, S. et al. 2012. Paramyxean–microsporidian co-infection in amphipods: Is the consensus that Microsporidia can feminise their hosts presumptive? – Int. J. Parasitol. 42: 683–691.

Silva, A. 2003. Morphometric variation among sardine (*Sardina pilchardus*) populations from the northeastern Atlantic and the western Mediterranean. – ICES J. Mar. Sci. 60: 1352– 1360.

Šimková, A., et al. 2000. Co-existence of nine gill ectoparasites (*Dactylogyrus*: Monogenea) parasitising the roach (*Rutilus rutilus* L.): history and present ecology. - Int. J. Parasitol. 30: 1077–1088.

Šimková, A., et al. 2004. Molecular phylogeny of congeneric monogenean parasites (*Dactylogyrus*): a case of intrahost speciation. – Evolution 58: 1001–1018.

Sitjà-Bobadilla, A. 2008. Living off a fish: a trade-off between parasites and the immune system. – Fish Shellfish Immunol. 25: 358–372.

Soler-Jiménez, L. C. and Fajer-Ávila, E. J. 2012. The microecology of dactylogyrids (Monogenea: Dactylogyridae) on the gills of wild spotted rose snapper *Lutjanus guttatus* (Lutjanidae) from Mazatlan Bay, Mexico. – Folia Parasitol. (Praha). 59: 53–58.

Stumbo, A. D. et al. 2012. Shoaling as an antiparasite defence in minnows (*Pimephales promelas*) exposed to trematode cercariae. – J. Anim. Ecol. 81: 1319–1326.

Teske, P. R. et al. 2021.The sardine run in southeastern Africa is a mass migration into an ecological trap. – Sci. Adv. 7: eabf4514.

Timi, J. T. and Poulin, R. 2003. Parasite community structure within and across host populations of a marine pelagic fish: How repeatable is it? – Int. J. Parasitol. 33: 1353– 1362.

Timi, J. T. et al. 2010. Similarity in parasite communities of the teleost fish *Pinguipes brasilianus* in the southwestern Atlantic: Infracommunities as a tool to detect geographical patterns. – Int. J. Parasitol. 40: 243–254.

Tinsley, R. C. et al. 2020. Heterogeneity in helminth infections: Factors influencing aggregation in a simple host-parasite system. – Parasitology 147: 65–77.

Tomnatik, V. E. 1990. The influence of water temperature on the sexual maturation of *Dactylogyrus vastator*. – Parazitologiya 24: 235–238.

van Zwieten, P. A. M. et al. 2002. Effects of inter-annual variability, seasonality and persistence on the perception of long-term trends in catch rates of the industrial pelagic purse-seine fishery of northern Lake Tanganyika (Burundi). – Fish. Res. 54: 329–348.

Vanhove, M. P. M. et al. 2021. From the Atlantic coast to Lake Tanganyika: Gill-infecting flatworms of freshwater pellonuline clupeid fishes in West and Central Africa, with description of eleven new species and key to *Kapentagyrus* (Monogenea, Dactylogyridae). – Animals 11: 3578.

Višnjić-Jeftić, Ž., et al. 2013. The geometric morphometrics and condition of Pontic shad, *Alosa immaculata* (Pisces: Clupeidae) migrants to the Danube River. - J. Nat. Hist. 47: 1121–1128.

Wang, C. et al. 2017. Zero-inflated hierarchical models for faecal egg counts to assess anthelmintic efficacy. – Vet. Parasitol. 235: 20–28.

Wickham, H. 2016. ggplot2: Elegant Graphics for Data Analysis. – Springer-Verlag.

Wickham, H. et al. 2019. Welcome to the Tidyverse. – J. Open Source Softw. 4: 1686.

Wilke, C. 2020. ggtext: Improved Text Rendering Support for “ggplot2”. R package version 0.1.1.

Wilson, A. B. et al. 2008.Marine incursion: the freshwater herring of Lake Tanganyika are the product of a marine invasion into west Africa. – PLoS One 3: e1979.

Windsor, D. A. 1998. Most of species on Earth are parasites. – Int J Parasitol 28: 1939–41.

Wood, C. L. et al. 2014. Fishing drives declines in fish parasite diversity and has variable effects on parasite abundance. – Ecology 95: 1929–1946.

Zhi, T. et al. 2018. Expression of immune-related genes of Nile tilapia *Oreochromis niloticus* after *Gyrodactylus cichlidarum* and *Cichlidogyrus sclerosus* infections demonstrating immunosupression in coinfection. – Fish Shellfish Immunol. 80: 397–404.

Zuur, A. F. et al. 2009. Zero-Truncated and Zero-Inflated Models for Count Data. – In: Mixed effects models and extensions in ecology with R. Statistics for Biology and Health. Springer, New York, NY, pp. 261–293.

