## Supplementary File for "Host lifestyle and parasite interspecific facilitation mediate co-infection in a species-poor host-parasite system"

**Supplementary methodology geomorphometrics**

To extract the shape information, full Procrustes fits of landmark data were performed and aligned by principal axes. Three classifiers (species, locality of origin, and subbasin) were imported for further analyses. Principal Component Analysis (PCA) was performed on the covariance matrix to visualise the shape variation. Highly deviating specimens, identified by the PCA plot, were excluded from the analysis. Regressions against the standard length of each specimen followed by a 10,000 replicate permutation test were performed on the first three individual PC axes. Due to the significant correlation between the standard length and PCA loadings/Procrustes distances (see Fig. S2), the final PCAs were performed on residuals which resulted from the regression analyses of Procrustes distances and standard length.

**Supplementary Figures**

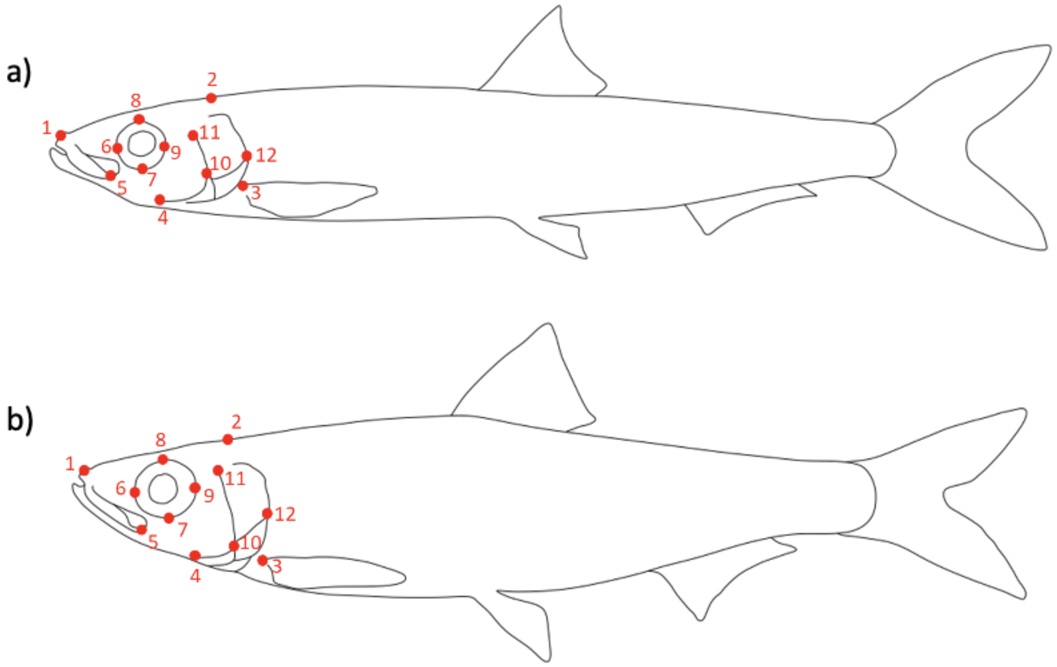


Fig. S1: Position of landmarks recovered for 2D digitisation of a) *Limnothrissa miodon* and b) *Stolothrissa tanganicae*.

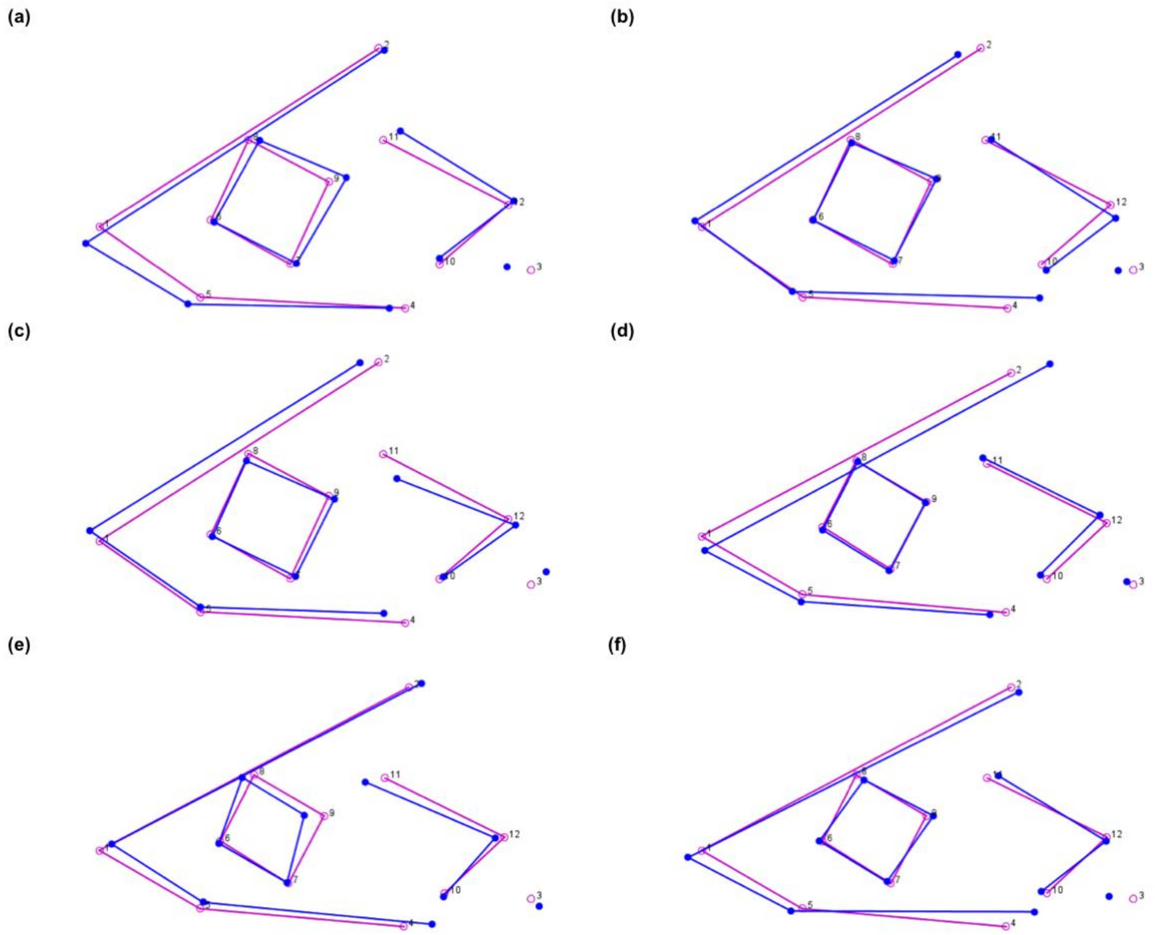


Fig. S2: Wireframes showing the shape variation in the head based on the coordinates of Principal Component Analyses of a) *Limnothrissa miodon,* PC1, b) *Limnothrissa miodon,* PC2, c) *Limnothrissa miodon,* PC3, d) *Stolothrissa tanganicae,* PC1, e) *Stolothrissa tanganicae,* PC2, f) *Stolothrissa tanganicae*, PC3. The target shape is presented in blue, the starting shape in pink.

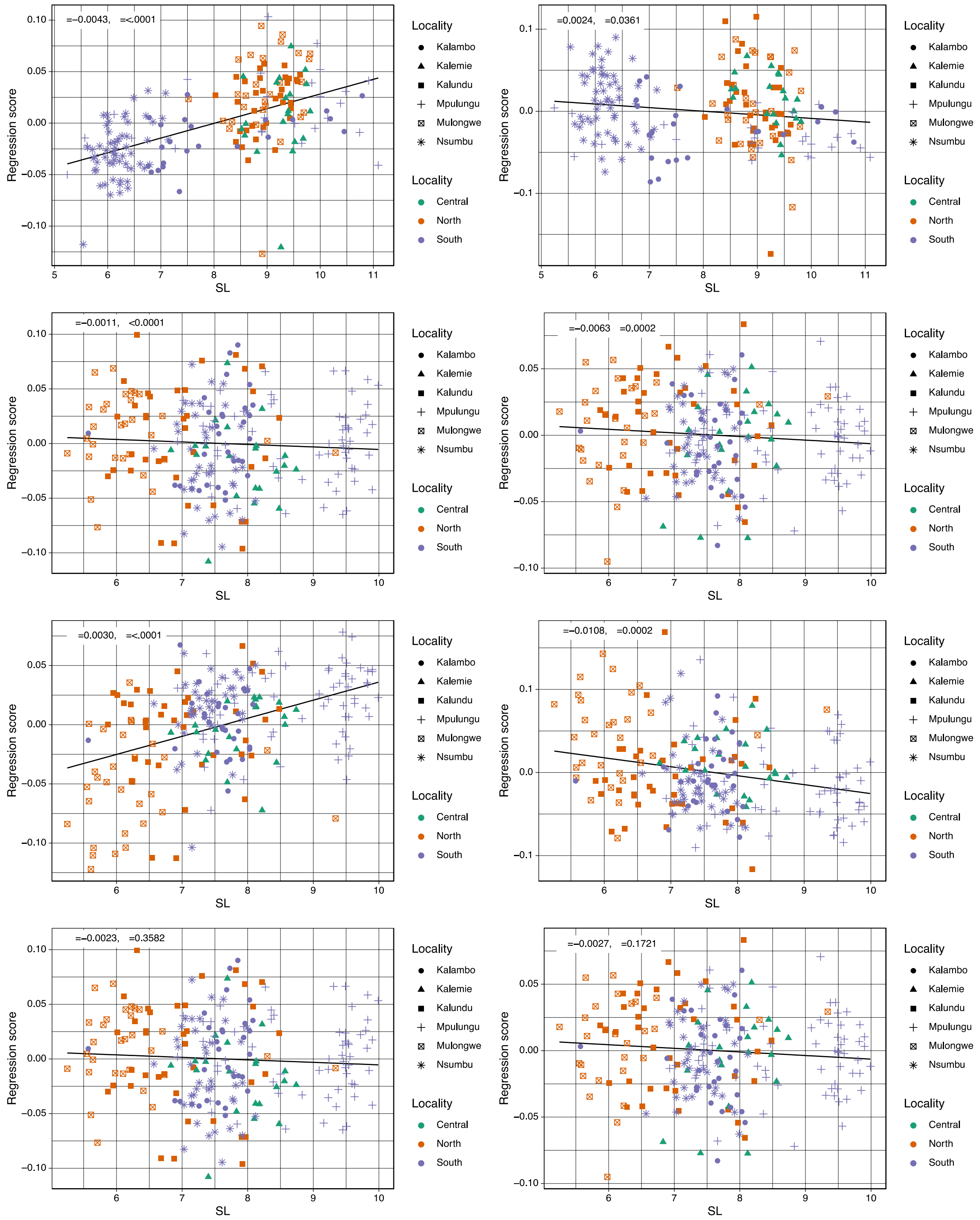


Fig. S3: Biplots showing results of regression analyses of the head across the sampling localities of a) Procrustes distances/coordinates and standard length (SL) of *Limnothrissa miodon*, b) Principal component scores of PC1 and SL of *Limnothrissa miodon*, c) Principal component scores of PC2 and SL of *Limnothrissa miodon*, d) Principal component scores of PC3 and SL of *Limnothrissa miodon*, e) Procrustes distances/coordinates and standard length

(SL) of *Stolothrissa tanganicae*, f) Principal component scores of PC1 and SL of *Stolothrissa tanganicae*, g) Principal component scores of PC2 and SL of *Stolothrissa tanganicae*, h) Principal component scores of PC3 and SL of *Stolothrissa tanganicae*. Regression coefficients and P-values are displayed in the left upper corner of each biplot.
